# Species Distribution, Antifungal Susceptibility, and Machine Learning-Based Prediction of Optimal Therapies Against *Candida* Species

**DOI:** 10.64898/2026.10.06.757060

**Authors:** Punit Tiwari, Amit Kumar Rai, Sirimani Varnika, Munesh Kumar Gupta, Ragini Tilak, Pradip Paik

## Abstract

Invasive candidiasis has become more common in recent decades, particularly within ICUs. Identifying specific *Candida* spp., and testing their sensitivity to antifungal drugs is crucial for effective treatment that helps healthcare providers to detect potential treatment challenges early. This study aimed to identify *Candida* spp. at their spp. level present in blood, assess their susceptibility to antifungal agents, and apply a random forest classifier to predict the best antifungal drug for a given species. The isolates were classified at the species complex level through MALDI-TOF MS. The strains’ susceptibility to antifungal agents such as AMB, FLZ, VOZ, ITZ, POZ, CAS, MFN, and ANF was assessed using CLSI guidelines. Out of the 354 *Candida* spp. isolated from blood, *C. utilis* was found to be the predominant species, followed by *C. tropicalis,* and *C. albicans*. This study demonstrated that MFN and ANF exhibited high potency, with an impressive MIC_90_ of 0.12 µg/mL against all isolates, surpassing CAS, which had a slightly higher overall MIC_90_ of 0.25 µg/mL. FLZ exhibited the highest MIC_90_ value among the azoles at 16 µg/mL, while VOZ, ITZ, and POZ all had MIC_90_ values of 0.5 µg/mL. AMB MIC_90_ values were 2 µg/mL against all the isolated *Candida* species. The Random Forest AI/ML model showed perfect overall prediction, with the actual and predicted best antifungal drugs matching for all fungal species. This study is particularly essential because rare *Candida* spp. becoming more common and exhibiting higher MICs to AMB and FLZ and required suitable remedies for controlling the infection.

**IMPORTANCE:** This study examines the incidence of common as well as rare *Candida* species and their susceptibility to different antifungal drugs. Here, we analyzed over 354 isolates of yeast from blood cultures of patients admitted to ICUs. Notably, this study reports the emergence of *C. utilis* in large numbers, alongside other rare yeast species in NICU. These findings highlight the increasing resistance against common as well as rare *Candida* species isolates from the northern site of India.

## INTRODUCTION

Invasive candidiasis has become more common in recent decades, particularly within Intensive Care Units (ICUs), and remains associated with a high rate of illness and death (1, 2). Neonates, immunocompromised patients, and patients with long-term ICU stays are a vulnerable group for *Candida* colonization and subsequent cause bloodstream infection (3, 4). According to various studies in the US, among hospital-associated bloodstream infections, candidemia is 4^th^ leading cause (3). Furthermore, there is also evidence of an increase in candidemia cases in India, Europe, and Australia (5). In terms of the relative frequency of different *Candida* spp. in bloodstream infections, it has altered in recent years (5, 6). The majority of invasive candidiasis, over 90%, is caused by five species: *C. albicans, C. tropicalis, C. parapsilosis, C. glabrata* (syn. *Nakaseomyces glabratus*)*, and C. krusei* (syn. *Pichia kudriavzevii*) (7). *C. albicans* used to be the predominant *Candida* spp. in clinical samples, but there is now a significant rise in infections from other *Candida* spp. (8)*. C. tropicalis* is the most frequently isolated *Candida* spp., especially in tropical climates, and it accounts for 3 to 66% of cases of candidemia (9).

It is necessary to identify the specific *Candida* spp. causing invasive candidiasis/BSIs to commence suitable and successful antifungal therapy (10). Conventional biochemical and morphological techniques frequently fail to identify rare or closely related yeasts and may misclassify them as more common species (11, 12). Among these, MALDI-TOF MS has emerged as a rapid, accurate, and cost-effective technique for identifying *Candida* spp, at the species or even subspecies level (13). Notably, MALDI-TOF MS has demonstrated high accuracy in the identification of rare and emerging *Candida* spp., which are often challenging to detect using traditional biochemical or morphological approaches (12).

Furthermore, antifungal susceptibility testing (AFST) using a standardized method is critical for guiding clinical decisions in managing fungal infections, tracking resistance, and comparing the efficacy of antifungal agents against these pathogenic yeasts (9, 14). The treatment of fungal infections is getting increasingly difficult as *Candida* spp. develop resistance to antifungal medicines (15). The sensitivity of various *Candida* spp. to antifungal drugs varies and can significantly influence the clinical outcomes. Due to the widespread use of antifungal drugs, multidrug resistance (MDR) has emerged in *Candida* species. Crucially, there is a non-uniform antifungal sensitivity profile across different species of *Candida*; precise identification coupled with routine susceptibility profiling is important to minimize clinical failure and increase patient outcomes.

Integrating antifungal susceptibility data with AI/ML models may help predict the best antifungal drug for a given species, thereby shortening treatment time and improving early clinical decision-making in ICU patients. In our study, we applied the Random Forest classifier as it effectively classifies the most appropriate antifungal drug based on the calculated Weighted MIC values of eight antifungal agents, handles complex nonlinear relationships among the MIC data, is less prone to overfitting than many individual classifiers, and provides reliable prediction accuracy.

In this retrospective study, carried out in a single facility in S.S. Hospital, Varanasi, India, over a 4-year period that covers a large population of northern states of India. Yeast isolated from blood samples underwent species identification and *in vitro* testing with eight antifungal drugs using the CLSI guidelines.

## MATERIALS AND METHODS

### Ethical Approval and Isolate Collection

This study received ethical clearance from the Institutional Review Board (Ref. no.-ECR/526/Inst/UP/2019). From January 2020 to December 2023, 354 non-duplicate strains of various *Candida* spp. were isolated from blood cultures of ICU patients admitted to S.S. Hospital (Varanasi, India). Prior to experimental evaluation, all collected isolates were stored at -80°C in 20% glycerol.

### Identification of *Candida* species

Species identification followed standard procedures, including colony morphology on cornmeal agar, chromogenic agar for color analysis (HiCrome *Candida* differential HiVeg agar, Himedia, Mumbai, India), morphology observation by Gram staining, germ tube formation, sugar assimilation, and sugar fermentation. Further identification of yeast isolates was done by MALDI-TOF MS (Bruker Daltonik GmbH, Bremen, Germany) as described by Ghosh et al. (2015), carried out at PGIMER Chandigarh (13). Prior to experimental testing, all isolates were sub-cultured on SDA for 24 hours under aerobic conditions at 37℃ to ensure the viability and purity of the isolated strains.

### Drugs and stock solutions

Eight antifungal agents commonly used in candidemia management were included for susceptibility testing: amphotericin B (AMB), fluconazole (FLZ), itraconazole (ITZ), voriconazole (VOZ), posaconazole (POZ), caspofungin (CAS), micafungin (MFN), and anidulafungin (ANF). AMB, POZ, ITZ, FLZ, ANF, and VOZ were obtained from Sigma-Aldrich (Germany), while CAS and MFN were procured from HiMedia (Mumbai, India). Stock solutions of AMB, POZ, ITZ, FLZ, ANF, and VOZ were prepared in dimethyl sulfoxide (DMSO), whereas CAS and MFN were dissolved in sterile distilled water, in accordance with CLSI recommendations. All stock solutions were stored at -80 °C until use. Working dilutions were subsequently prepared using RPMI-1640 broth medium according to the CLSI protocols.

### Antifungal Susceptibility Assay

AFST of *Candida* isolates was determined using the CLSI M27-A4 broth microdilution method (16). Antifungal drugs were serially diluted (0.03-64µg/ml) into a 96-well flat-bottom microtiter plate. Fungal inocula were generated from 24 h cultures grown on SDA at 37℃ in sterile 0.85% NaCl solution. The suspended cell density was adjusted to 0.5 McFarland standard. This was followed by subsequent dilution in RPMI 1640 medium to achieve a final inoculum density of roughly 5.0 × 10² to 2.5 × 10³ cells/mL. After a 24 h incubation at 35℃, MICs were interpreted according to the CLSI guidelines (17). For quality control, *C. krusei* ATCC 6258 and *C. parapsilosis* ATCC 22019 served as reference strains.

### Weighted MIC Calculation and Random Forest Classification Analysis

Raw antifungal susceptibility data were obtained from clinical blood-culture isolates. For each fungus-drug pair, minimum inhibitory concentration (MIC) distributions (MIC concentrations and counts of isolates inhibited at each concentration) were available. The MIC distribution for each fungus-drug combination into a single representative value, a weighted MIC was calculated as the count-weighted mean of MIC concentrations:

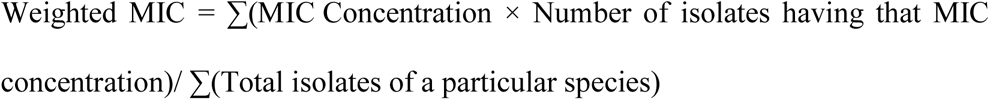

Weighted MIC values for eight antifungal agents were computed for each fungal species and compiled into a tabular dataset. Rows represent fungal species and columns represent the weighted MIC for each drug. The assembled table was exported to Excel for downstream analysis.

The best drug assignment for lower MIC indicates greater antifungal activity; the antifungal agent with the minimum weighted MIC for each fungal species was identified using a row-wise minimum (idxmin) operation.

### Software and libraries

The analysis and modeling was done in Python(3.14) using pandas for data manipulation, matplotlib and seaborn for visualization, and scikit-learn for model training and evaluation (18). Since scikit-learn classifiers need numerical target labels, LabelEncoder was used to convert categorical drug names in best_drug to integer labels, keeping a map from drug names to numeric labels to translate predictions back to drug names (19, 20). The Random Forest Classifier was chosen for its robustness to correlated features and its ability to capture complex, non -linear decision boundaries. The model was configured with 300 estimators (n_estimators = 300) to reduce prediction variance and improve the stability and reliability of the results (21).

### Cross-validation and evaluation

For model validation, Leave -One-Out Cross-Validation (LOOCV) was employed to provide a robust estimate of the model’s generalization performance, particularly given the limited number of fungal species. In this approach, one fungal species was excluded as the test sample while the model was trained using all remaining species. The procedure was repeated until each fungal species had served as the test sample exactly once. For each fold, the trained model generated a predicted class labels were subsequently converted back to their corresponding drug names and aggregated for evaluation. Model performance was then assessed using standard multiclass classification metrics based on the complete set of LOOCV predictions (22). Accuracy represented the proportion of fungal species correctly classified, precision indicated the proportion of predicted drug classes that were correct, recall measured the proportion of actual drug classes correctly identified, and the F1 score represented the harmonic mean of precision and recall. These metrics provided a comprehensive assessment of the classifier’s predictive performance across the different drug classes (23).

### Model training, final predictions and visualization

After cross-validation, the Random Forest classifier was retrained on the entire dataset (all fungal species) using the same hyperparameters. The final model was applied to the dataset to produce predicted Best_Drug labels for each fungal species. A results table was produced containing fungal species name, actual Best_Drug (derived from minimum weighted MIC), and model-predicted Best_Drug. Weighted MIC distributions per drug and species were visualized using boxplots to inspect feature distributions and identify potential outliers. Classification metrics (accuracy, precision, recall, F1) were visualized as bar charts to facilitate interpretation.

### Data management and statistical analysis

The datasets were independently verified with Microsoft Excel (Redmond, Washington, USA) to eliminate possible errors. For each species, MIC ranges, MIC₅₀, and MIC₉₀ were calculated against the antifungal agent tested. Species-level MIC distributions were summarized to highlight interspecies variation in susceptibility. While established CLSI clinical breakpoints and epidemiological cutoff values (ECVs) were used where available, susceptibility interpretation for rare species, such as *C. utilis*, relied primarily on MIC distributions and comparisons with other *Candida* species, given the lack of species-specific breakpoints. A Chi-square test of independence was performed to evaluate the distribution of *Candida* spp. by gender. A two-way ANOVA with Tukey’s multiple-comparison test was performed to assess variations in MIC_50_ and MIC_90_ values across various antifungal drugs. *P-values* < 0.05 were considered statistically significant. Data were analyzed and visualized using GraphPad Prism v.8.4.2 (GraphPad Software, San Diego, CA, USA).

## RESULTS

### Species distribution

In a duration of 4 years, a total of 354 non-duplicate *Candida* spp. were isolated, of which 223 were male, and 131 were female (Table **1**). Major predisposing risk factors were septic shock (78.81%) and respiratory disease (75.58%), followed by vascular access (52.22%), preterm (44.35%), low birth weight (38.13%), and broad-spectrum antibiotics (33.49%), as shown in Table **2**. In this study, 91.52% of the total isolates were non-albicans *Candida*, whereas C. *albicans* accounted for only 8.47%. Commonly encountered *Candida* spp. showed specific colored colonies on CHROMagar, and morphological features such as hyphae, pseudohyphae, blastospores, and ascospores of some *Candida* spp. were visualized on corn meal agar supplemented with Tween 80, as shown in Figures 1 and 2, respectively.

**Figure 1.**
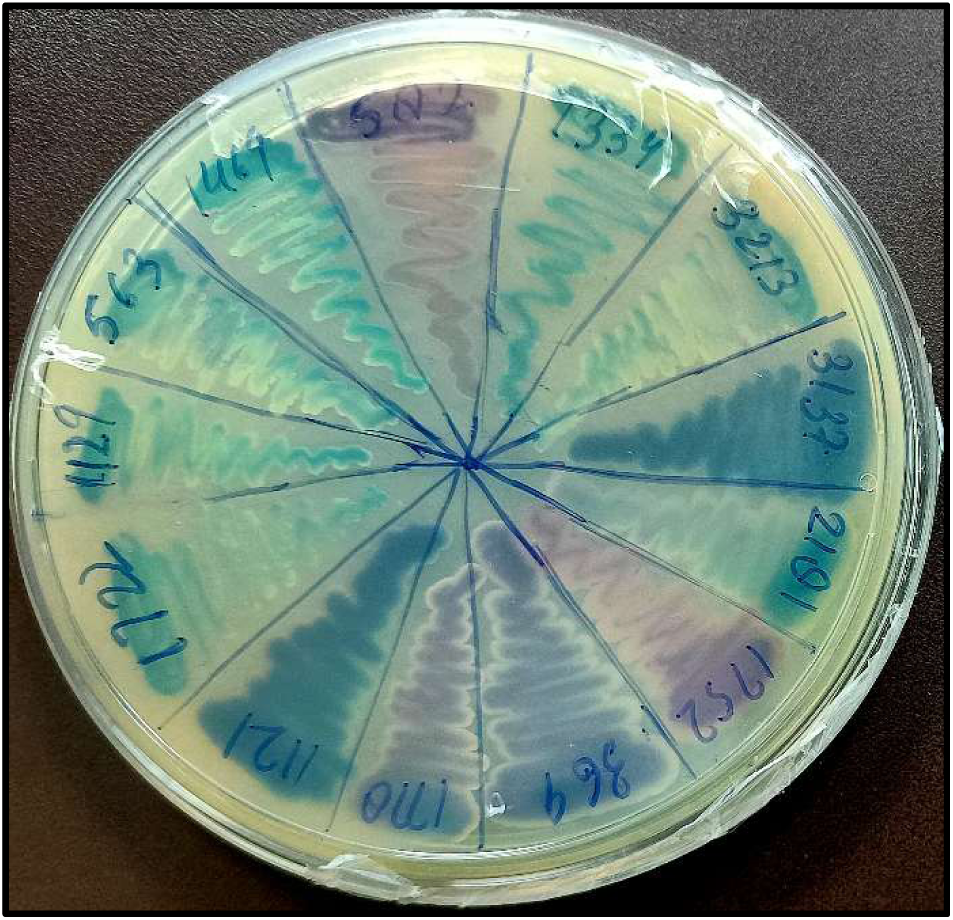
Chrom agar images of *Candida* reveal color-based identification of isolated species.

**Figure 2.**
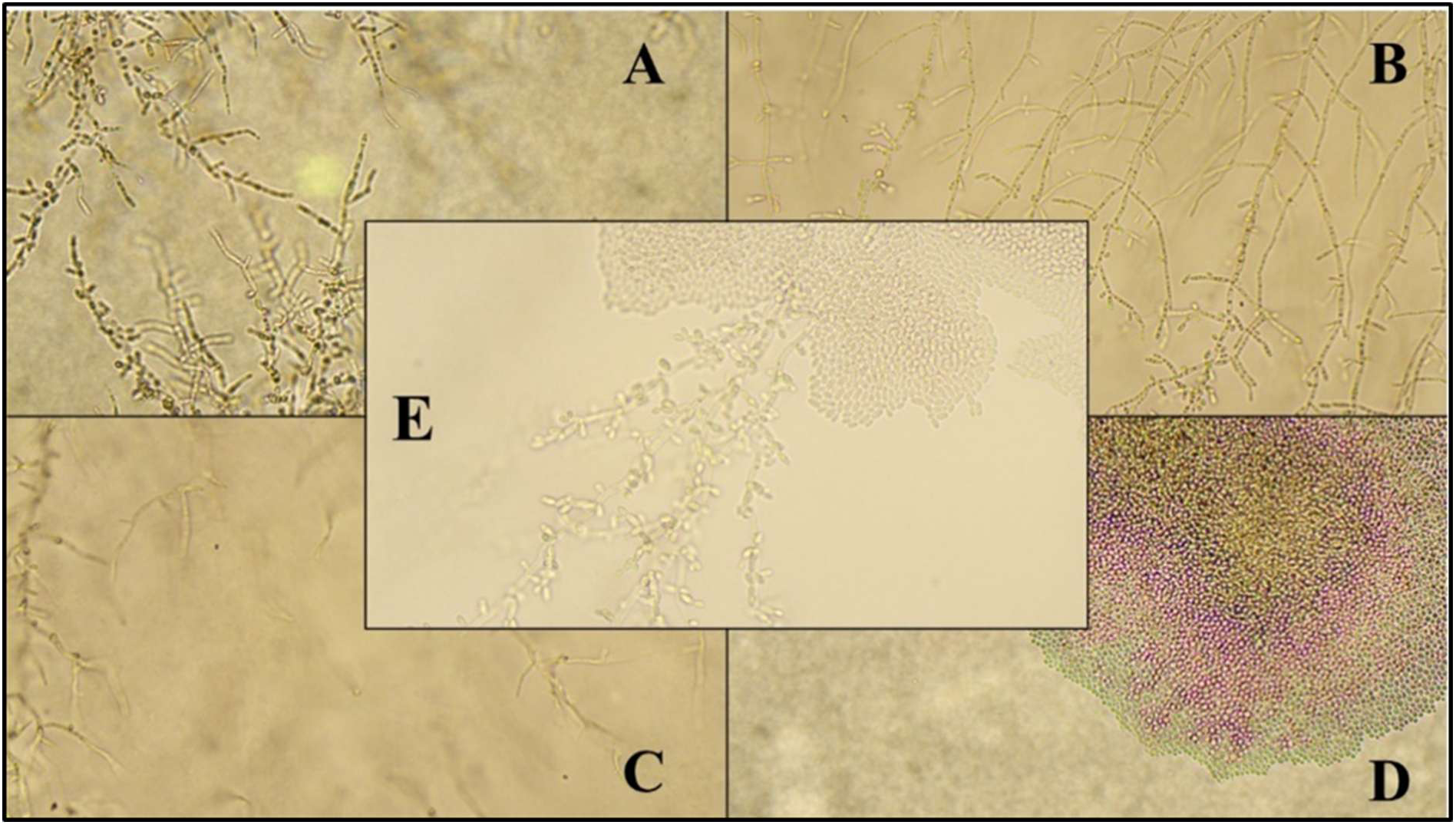
Morphology of Candida spp. (A) C. albicans; (B) C. tropicalis; (C) C. parapsilopsis; (D) *C. auris; (E) C. utilis* on Corn meal agar (40x).

**Table 1.** Number and Prevalence (%) of *Candida* spp. in blood and their distribution by patient gender.

| S. No. | Species | No. of<br>Isolates | Percentage | Male | Female |
| --- | --- | --- | --- | --- | --- |
| 1 | *Candida utilis* (*Cyberlindnera jadinii*) | 146 | 41.24 | 92 | 54 |
| 2 | *Candida albicans* | 30 | 8.47 | 19 | 11 |
| 3 | *Candida tropicalis* | 92 | 25.98 | 52 | 40 |
| 4 | *Candida parapsilosis* | 4 | 1.12 | 3 | 1 |
| 5 | *Candida krusei* (*Pichia kudriavzevii*) | 7 | 1.97 | 5 | 2 |
| 6 | *Candida glabrata* (*Nakaseomyces glabratus*) | 6 | 1.69 | 3 | 3 |
| 7 | *Candida pelliculosa*<br>(*Wickerhamomyces anomalus*) | 39 | 11.0 | 27 | 12 |
| 8 | *Candida auris* (*Candidozyma auris*) | 17 | 4.8 | 13 | 4 |
| 9 | *Kodamaea ohmeri* | 7 | 1.97 | 4 | 3 |
| 10 | *Candida orthopsilosis* | 1 | 0.282 | 1 | 0 |
| 11 | *Candida metapsilosis* | 1 | 0.282 | 1 | 0 |
| 12 | *Candida rugosa* | 4 | 1.12 | 3 | 1 |
|  | Total | 354 | 99.924 | 223 | 131 |

**Table 2.** Risk factors among the patients having suspected bloodstream *Candida* infection.

| Risk factor | Percentage of patients |
| --- | --- |
| Preterm | 44.35 |
| Low birth weight | 38.13 |
| Broad-spectrum antibiotics | 33.49 |
| Ventilator support | 10.16 |
| Vascular accesses | 52.22 |
| Gestational period | 14.12 |
| Septic shock | 78.81 |
| Respiratory diseases | 75.58 |
| Congenital diseases | 8.4 |
| Others<sup>a</sup> | 0.5 |

*Note:* <sup>a</sup> Feed intolerance, Acute and chronic renal failure, post-operative patients

All 354 isolated strains were further identified by MALDI-TOF MS. The most frequently isolated species was *C. utilis* (146 isolates, 41.24%), followed by *C. tropicalis* (92 isolates, 25.98%) and *C. albicans* (30 isolates, 8.47%). Other species included *C. pelliculosa* (11.01%), *C. auris* (17, 4.80%), *K. ohmeri* (7, 1.97%), *C. krusei* (7, 1.97%), *C. glabrata* (6, 1.69%), *C. parapsilosis* (4, 1.12%), *C. rugosa* (4, 1.12%), and single isolates each of *C. orthopsilosis* and *C. metapsilosis* (0.28%), as shown in Table **1**. All isolates of *C. utilis*, *C. pelliculosa*, *K. ohmeri,* and *C. orthopsilosis* were recovered from the NICU.

### Antifungal susceptibility tests

MICs of eight antifungal drugs, namely AMB, FLZ, VOZ, ITZ, POZ, CAS, MFN, and ANF, were determined for various *Candida* spp., and findings suggest variability in MIC values among different *Candida* isolates, as shown in Table **3**. In this study, among the azoles, only FLZ had the highest overall MIC_90_ value (16 µg/mL), whereas VOZ, ITZ, and POZ had overall MIC_90_ values of 0.5 µg/mL, as shown in Table **3**. The MIC range for FLZ was 0.12-≥64µg/mL, for VOZ it was ≤0.06-8µg/mL, and for both ITZ and POZ, the MIC range was ≤0.06-2µg/mL. Of the isolates studied, six isolates of *C. tropicalis*, one isolate of both *C. albicans* and *C. parapsilosis* show MIC value ≥ 8µg/mL against FLZ (Table **SI** of supplementary material). Rare *Candida* spp. like *C. utilis*, *C. pelliculosa*, *C. rugose*, *C. auris*, and *K. ohmeri* show MIC_50_ and MIC_90_ 1,4; 8,16; 32,64; 16,64; 4,8; 4,4; respectively for FLZ (Table **3**). For VOZ, ITZ, and POZ, MIC_90_ values for all isolated *Candida* spp. were the same, which is 0.5µg/mL. AMB MIC_90_ value was greater (≥4 µg/mL) in *C. utilis, C. parapsilosis, C. krusei*, and *C. auris* (Table **3**). All echinocandin drugs show good activity against most *Candida* isolates. On the basis of MIC_90_ value, CAS (0.5µg/mL) had a higher overall MIC_90_ value than MFN (0.12µg/mL) and ANF (0.12µg/mL) (Table **3**). In terms of overall MIC_90_ value, MFN and ANF were the most active drugs against all isolates (0.12 µg/mL) (Table **3**).

**Table 3.**
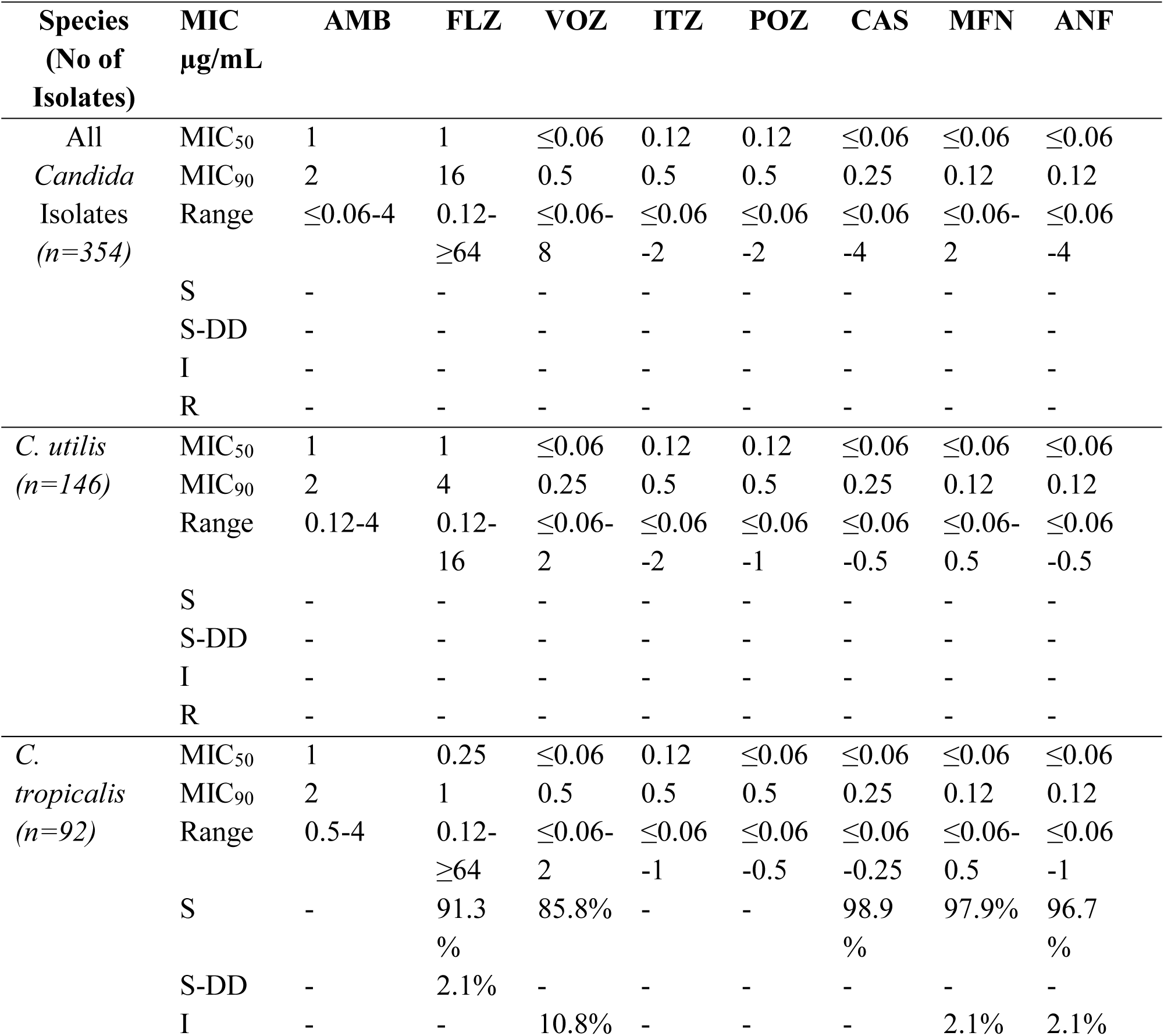
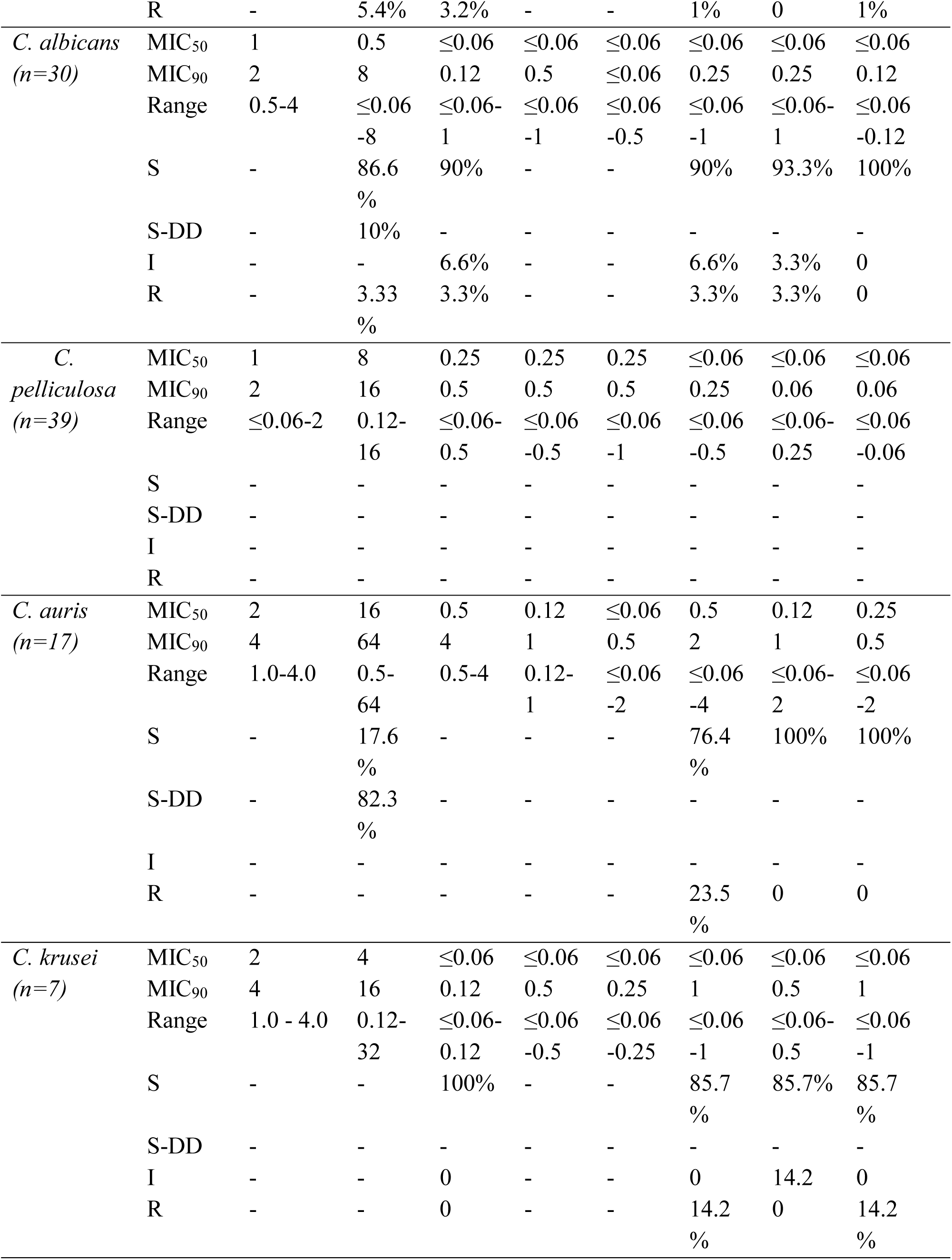
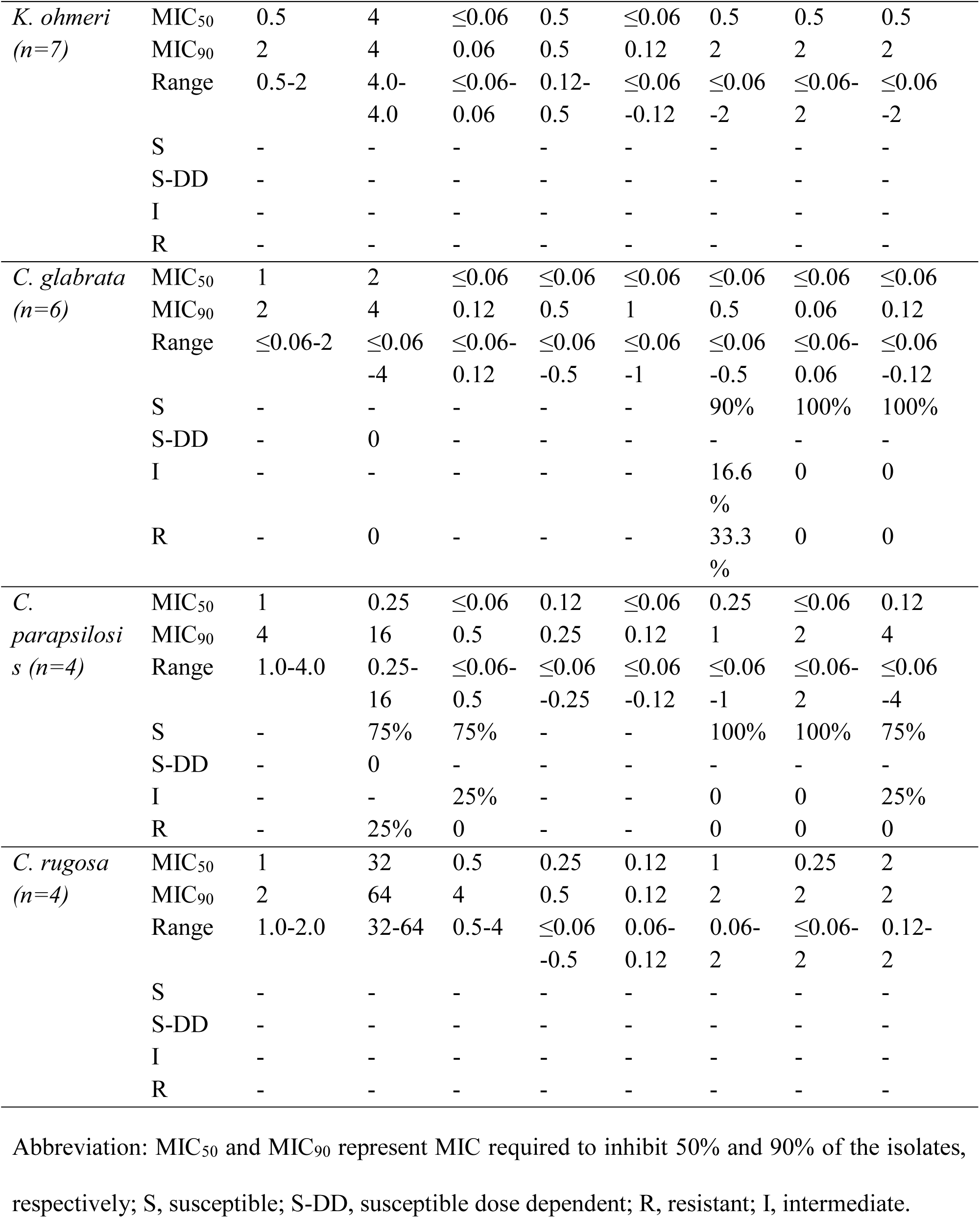
Antifungal Susceptibility data for *Candida* bloodstream isolates.

In FLZ, the resistance rate was observed 5.4% in *C. tropicalis*, 3.33% in *C. albicans* (3.33%), and 25% in *C. parapsilosis* (Table **3**). For VOZ, the resistance rate was observed in 3.2% in *C. tropicalis* and 3.33% in *C. albicans*. In CAS, 90% of isolates of *C. albicans*, 98.9% of isolates of *C. tropicalis*,76.4% of isolates of *C. auris*, 90% of isolates of *C. glabrata*, 100% of isolates of *C. parapsilosis,* and 85.7% of isolates of *C. krusei* were sensitive. For MFN, only *C. albicans* showed resistance of 3.3%, whereas all isolates of *C. tropicalis*, *C auris*, *C. krusei*, *C. glabrata*, and *C. parapsilosis* are 100% sensitive (Table **3**). In ANF, *C. tropicalis* and *C. krusei* isolates showed resistance of 1% and 14.2%, respectively, and other *Candida* spp., such as *C. tropicalis*, *C auris*, *C. krusei*, and *C. glabrata*, are 100% sensitive to ANF (Table **3**).

### Random forest classification

From Table 4, Random forest model predicted the best drug for particular *Candida* species. MFN was predicted for *C. tropicalis*, *C. utilis*, and *C. glabrata*; VOZ was predicted for *C. krusei*, *K. ohmeri*, *C. orthopsilosis*, and *C. metapsilosis*; ANI was predicted for *C. albicans* and *C. pelliculosa*; and POZ was predicted for *parapsilosis*, *C. auris*, and *C. rugosa*. The predicted drugs were identical to the actual best drugs across all 12 *Candida* species. The Random Forest classifier achieved an accuracy of 0.667 (66.7%), a precision score of 0.567 (56.7%), a recall score of 0.667 (66.7%), and F1 score of 0.604 (60.4%) as shown in fig. 5.

**Table 4.**
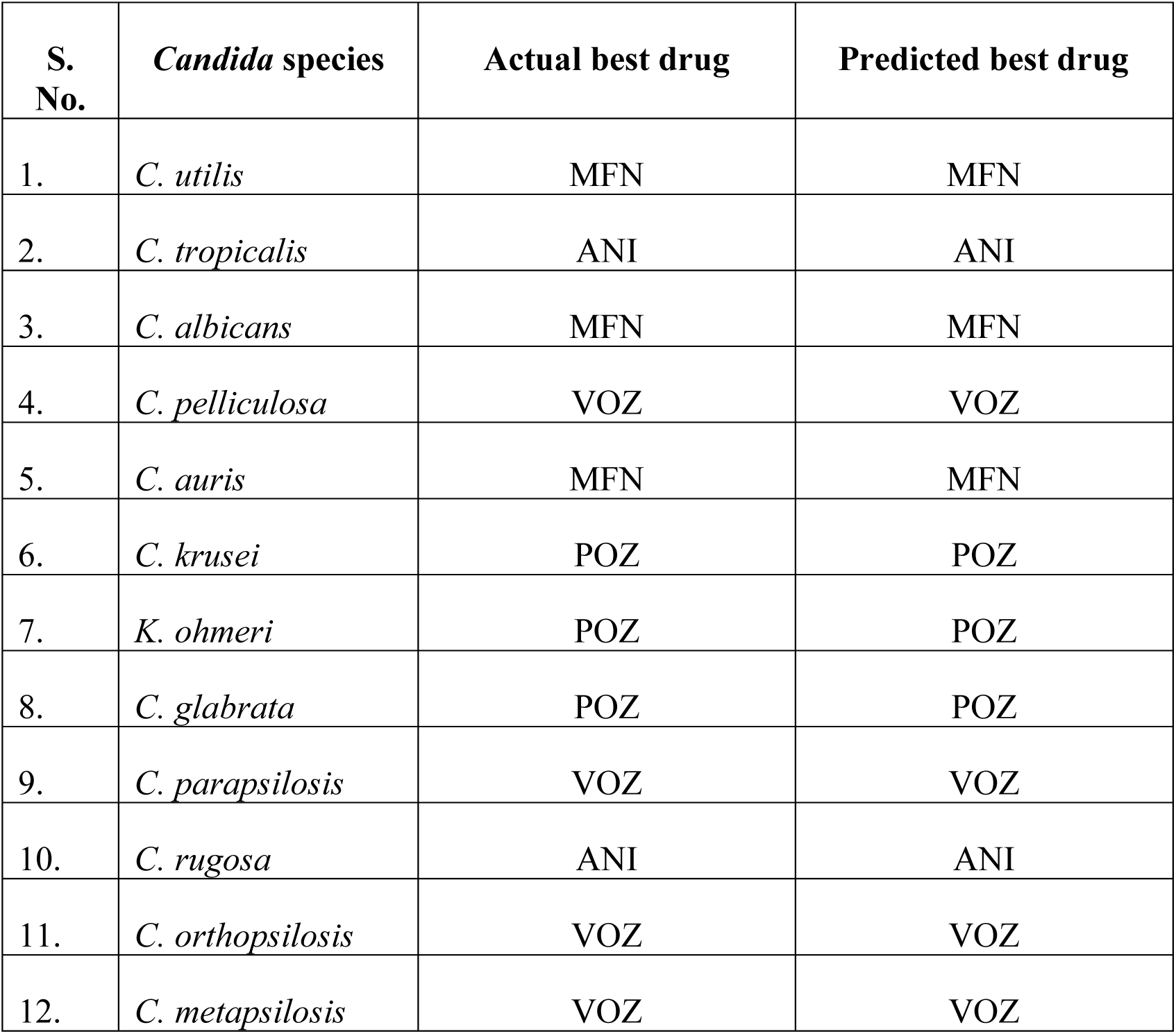
Comparison of actual and predicted best antifungal drugs for fungal species using the Random Forest classifier.

### Statistical analysis

There is no significant association between gender and the specific *Candida* species isolated. There is no significant difference between the MIC_50_ and MIC_90_ values for any specific drug. This indicates that isolates have highly uniform and homogeneous susceptibility profiles and that the drug’s behavior is highly predictable. Because MIC_90_ is not significantly higher than your MIC_50_, this strongly implies that no hidden, resistant subpopulations or non-wild-type (NWT) mutants are emerging in the sample group (Fig. **3**).

**Figure 3.**
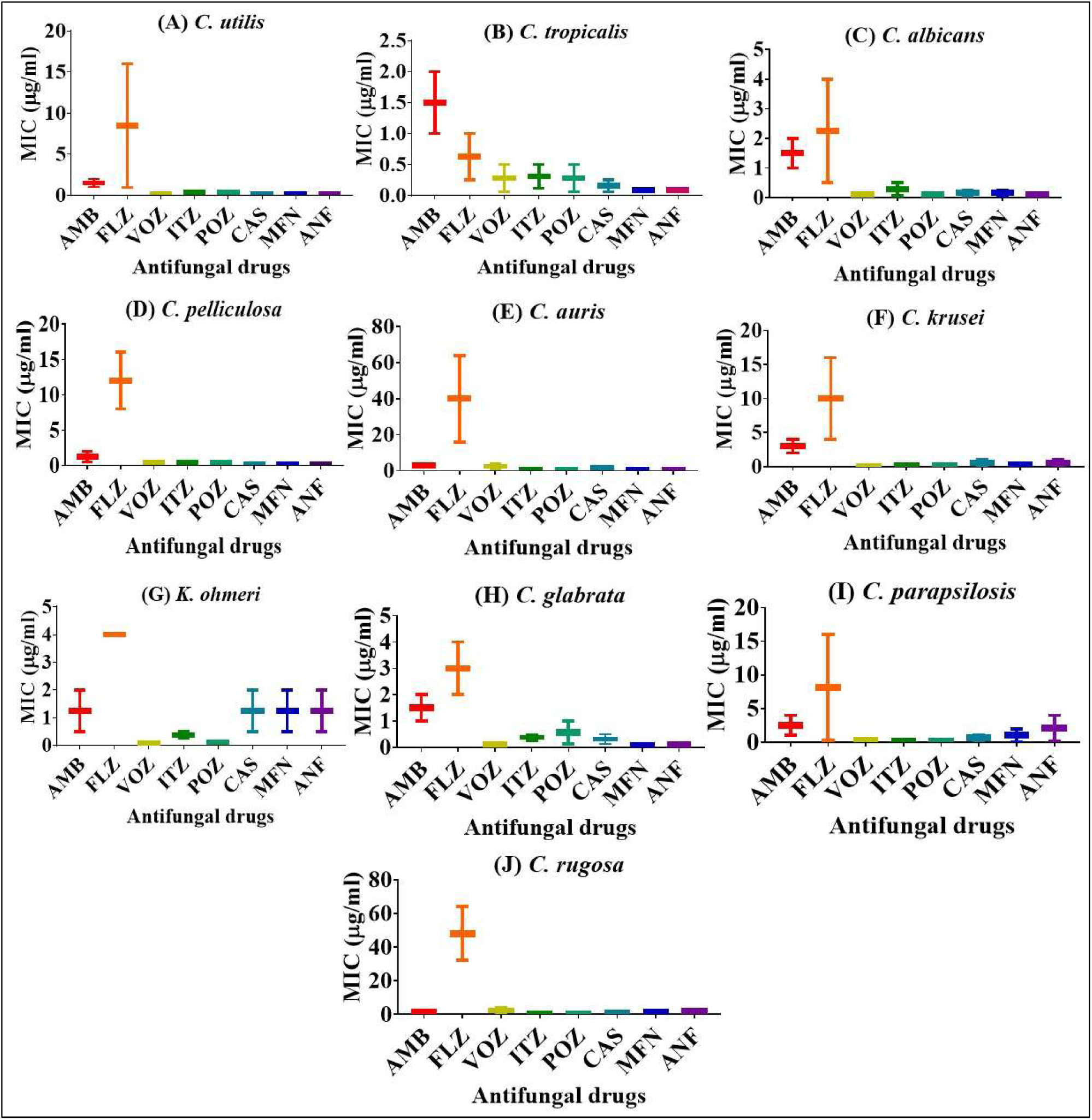
Whisker plot showing the minimum inhibitory concentration profiles (MIC_50_ and MIC_90_) of eight antifungal drugs (amphotericin B [AMB], fluconazole [FLZ], voriconazole [VOZ], itraconazole [ITZ], posaconazole [POZ], caspofungin [CAS], micafungin [MFN], and anidulafungin [ANF]) against isolated *Candida* species.

## DISCUSSION

Surveillance programs have investigated invasive *Candida* infections to identify the types of species involved and their response to antifungal treatments (24). Out of 354 *Candida* isolates we collected, *C. utilis* was found to be the predominant species, followed by *C. tropicalis, C. albicans,* C. *pelliculosa,* and *C. auris,* which together accounted for more than 90% of candidemia cases during 2020-2023. Similar trends in species distribution were reported, showing dominance of non-albicans *Candida* species in the Indian population (25). This study also revealed that 65.81% of the total *Candida* isolates from blood samples were obtained from the NICU. All uncommon *Candida* species and related yeast-like *C. utilis*, *C. pelliculosa*, *K. ohmeri*, and *C. orthopsilosis* were recovered from the NICU. This highlights the importance of speciation of these rare pathogens by advanced techniques like MALDI-TOF MS.

In the past, *C. albicans* was recognized as the main etiological agent of candidiasis. However, there has been a shift in the most common cause of candidemia from *C. albicans* to non-albicans *Candida* spp. (26). Bloodstream infections caused by non-albicans *Candida* spp. have increased significantly (27). In contrast to *C. albicans*, non-albicans *Candida* spp. display diverse responses to antifungal treatment. This difference could be associated with various virulence factors, including heightened production of proteinase and phospholipase enzymes (28). In this study, a high prevalence of NAC candidemia was observed, but there was a sudden increase in *C. utilis* candidemia cases due to an outbreak of *C. utilis* in the NICU of S.S. Hospital, Varanasi, Uttar Pradesh, India, during 2020 to 2023. From various environmental fomites and healthcare workers, swabs were taken and cultured to determine the probable sources of *C. utilis* in the hospital, but no isolates were recovered after culture. Although the telomorphic stage (*Cyberlindnera jadinii*) of *C. utilis* is widely utilized in the food and beverage industry, this uncommon yeast has also been reported as an opportunistic human pathogen previously in neonates (29, 30).

In this study, all isolates of *C. utilis*, *C. pelliculosa*, *K. ohmeri*, and *C. orthopsilosis* were recovered from the NICU, and very low birth weight, preterm birth, and septic shock were the main predisposing factors for neonatal candidemia (Table **2**). Other major risk factors that have been reported earlier for neonatal candidemia include prolonged labor, premature rupture of membranes, medical instrumentation during delivery, the use of antibiotics, maternal vaginal *Candida* colonization, and the extended use of indwelling catheters (31, 32). This diagnostic overlap highlights the critical importance of species identification to guide proper treatment in case of neonatal candidemia.

Various antifungal medications, including azoles (such as FLZ, ITZ, POZ, and VOZ), polyenes (such as AMB), and echinocandins (such as MFN, ANF, and CAS), are used to treat *Candida* infections. FLZ is commonly used to treat a broad spectrum of fungal and yeast infections (33). In this study, most *Candida* strains exhibit fluconazole MIC values of ≤4 µg/ml, except for specific strains: six of *C. tropicalis*, seventeen of *C. utilis*, sixteen of *C. auris,* twenty of *C. pelliculosa*, three of *C. rugosa*, three of *C. krusei*, one of *C. albicans,* and one of *C. parapsilosis,* this likely due to extended and or recurrent antifungal therapies. In this study, FLZ resistance rates were 3.33% for *C. albicans*, 5.4% for *C. tropicalis* and 25% for *C. parapsilosis*. These findings are comparable with published data, which indicate resistance rates of 2.3-5.8% for *C. albicans* and 6.2-9.8% for *C. tropicalis* and more than 30% for *C. parapsilosis* (26, 34–37). *C. glabrata* showed greater resistance rates (7.7-11.9%) compared to other *Candida* spp. against FLZ (38–40). However, in this study, all tested *C. glabrata* strains were susceptible to FLZ, with none showing resistance. Recent research assessing azole susceptibility across different *Candida* spp. revealed the lowest FLZ efficacy (41–44). This research also showed FLZ to be the least effective among the azoles, with a high MIC_90_ value for various *Candida* spp. including *C. pelliculosa* (MIC_90_ 16µg/mL), *C. auris* (MIC_90_ 64µg/mL), *C. rugosa* (MIC_90_ 64µg/mL), and *C. utilis* (MIC_90_ 4µg/mL). Heatmap further demonstrates this trend, where elevated MIC_50_ and MIC_90_ values suggest the emergence of FLZ resistance among the tested *Candida* species (Fig. **4**). In a previous study, VOZ, ITZ, and POZ from the azole group exhibited heightened efficacy against various *Candida* spp., displaying lower MIC_90_ values of 0.5µg/mL compared to FLZ (26, 44, 45). Interestingly, this current study yielded identical results, with an overall MIC_90_ of 0.5 µg/mL for these antifungal agents, except for *C. auris*, which had an MIC_90_ of 1µg/mL. Based on the CLSI M59 document, both the isolates of *C. orthopsilosis* and *C. metapsilosis* are wild-type strains for all classes of drugs used in this study (Table **SI** of supplementary material).

**Figure 4.**
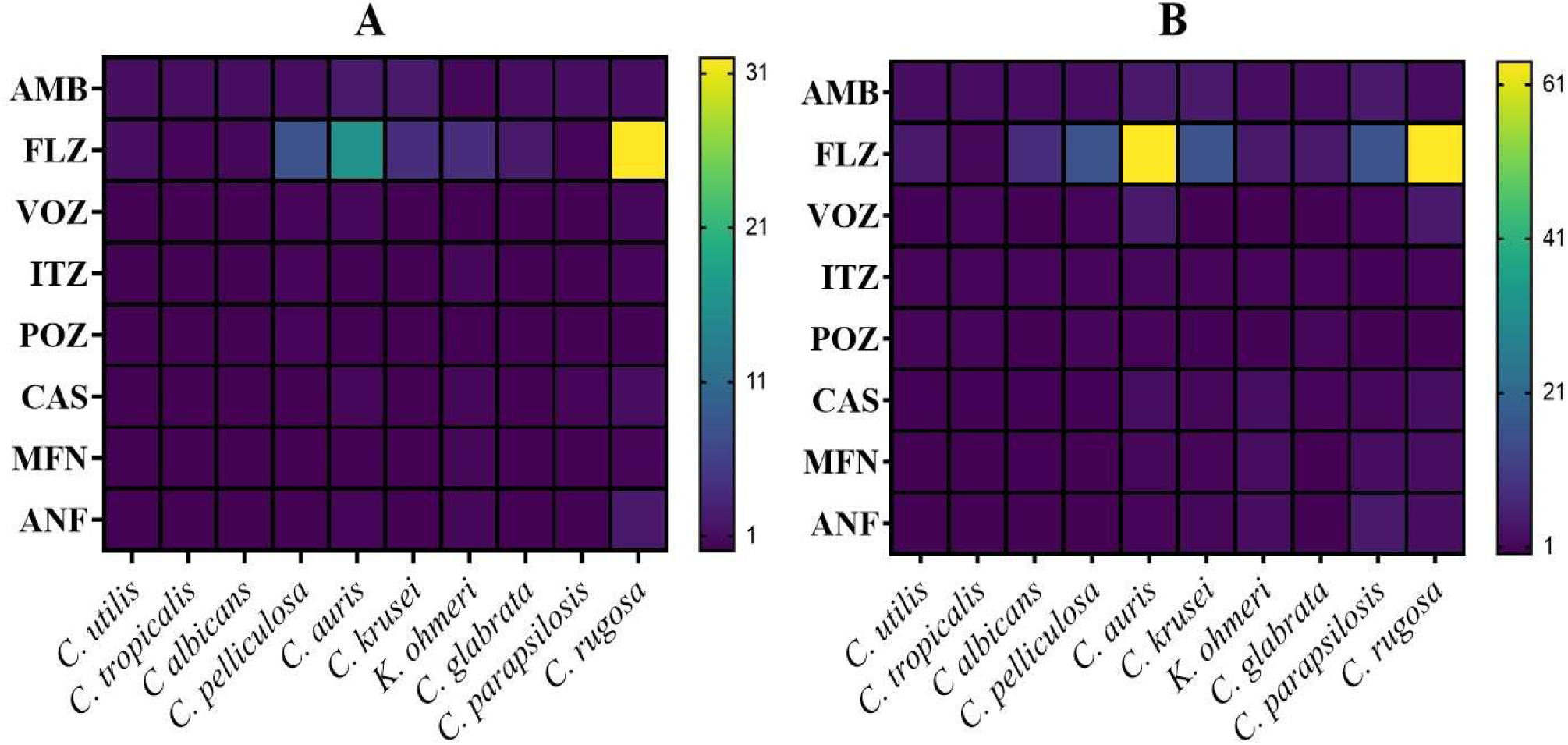
Heatmap of (A) MIC_50_ and (B) MIC_90_ values (µg/mL) of various antifungal agents against different *Candida* species. Bright yellow boxes indicate elevated (MIC_50_ & MIC_90_) values, notably demonstrating high fluconazole (FLZ) resistance in *C. auris* and *C. rugosa*.

**Figure 5.**
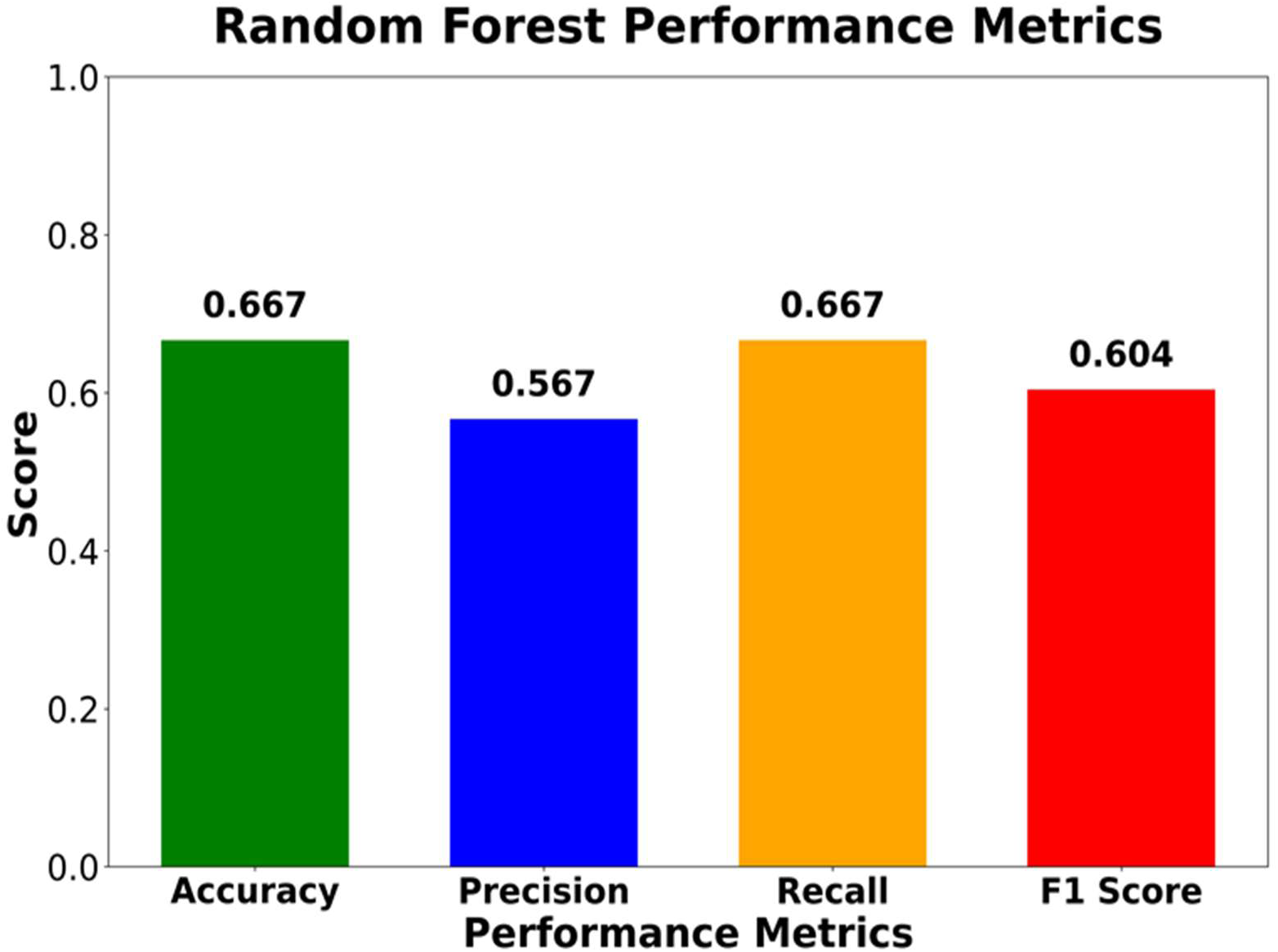
Performance Metrics of Random Forest, where the green bar indicates the accuracy score of 0.667, the blue bar indicates the precision of 0.567, the orange bar indicates the recall metrics of 0.667, and the red bar indicates the F1 score of 0.604.

For decades, AMB has been considered the most reliable and effective drug against severe and resilient fungal infection, despite its toxicity (46, 47). AMB resistance has been lower than that of azole antifungals among *Candida* spp., including *C. albicans*, *C. glabrata, C. parapsilosis, C. tropicalis, C. kefyr, and C. krusei* isolates (10). AMB lacks a specific breakpoint because it benefits the immune system and enhances the body’s defenses against fungal infections (48). In our investigation, an elevated AMB MIC_90_ value (≥4 µg/mL) was observed in *C. auris*, *C. parapsilosis*, and *C. krusei*, echoing similar low-susceptibility trends found in earlier investigations (49–52) (Table **3**). As shown in Table 3, the MIC range of AMB (≤0.06 to 4µg/mL) against *Candida* spp. was comparable to the previous study (53)

In this study, all three echinocandins had MICs of ≤2 μg/mL, covering 99.43% of *Candida* spp. isolates without distinction among species groups, a similar report was also published by Pfaller et al. (2008) (54). According to the CLSI species-specific susceptibility breakpoints, the isolates evaluated in this study demonstrated varied susceptibility profiles across the three echinocandins. For CAS, susceptibility rates were highest in *C. parapsilosis* (100%), followed by *C. tropicalis* (98.9%), *C. albicans* (90%), *C. glabrata* (90%), *C. krusei* (85.7%), and *C. auris* (76.4%). MFN exhibited 100% susceptibility against *C. glabrata*, *C. auris*, and *C. parapsilosis*, with slightly lower susceptibility rates observed for *C. tropicalis* (97.9%), *C. albicans* (93.3%), and *C. krusei* (85.7%). Lastly, susceptibility to ANF was 100% for *C. albicans*, *C. glabrata*, and *C. auris*, followed by *C. tropicalis* (96.7%), *C. krusei* (85.7%), and *C. parapsilosis* (75%) (55). These findings underscore species-specific variations in antifungal susceptibility, particularly highlighting heightened resistance patterns in certain *Candida* spp. to specific antifungal agents, providing valuable insights for clinical management and future research.

The Random Forest classifier successfully predicted the best antifungal drug for all fungal isolates tested. The predicted drugs were identical to the actual best drugs across 12 *Candida* species, indicating complete concordance between observed and model-predicted outcomes. This means the model performed perfectly on the displayed dataset, with no mismatches between observed and predicted drug choices.

The performance metrics of the Random Forest classifier indicate a moderate predictive performance for identifying the best antifungal drug. The model achieved an accuracy of 0.667 (66.7%), indicating that approximately two-thirds of the predictions were correctly classified. The precision score of 0.567 (56.7%) suggests that when the model predicted a particular drug as the best drug, about 56.7% of those predictions were correct. The recall score of 0.667 (66.7%) indicates that the model successfully identified approximately 66.7% of the actual best-drug classes. The F1 score of 0.604 (60.4%), which provides a balance between precision and recall, demonstrates moderate overall classification effectiveness. The higher recall compared with precision suggests that the model is relatively better at identifying the appropriate drug classes than at ensuring that every predicted drug is correct. Overall, these results demonstrate that the Random Forest model has moderate discriminative capability, but the relatively lower precision indicates that further improvement through additional data, feature selection, class balancing, or model optimization may be beneficial.

In conclusion, there is a crucial need to promptly and accurately identify *Candida* spp. and determine their MICs against different antifungal agents for appropriate antifungal therapy. MALDI-TOF MS was an important, reliable, and rapid tool for the identification of common as well as rare bloodstream yeasts. This is particularly essential because rare *Candida* spp. are becoming more common and exhibit higher MICs to AMB and FLZ. Notably, we have also reported a significant outbreak of *C. utilis* observed in the NICU over 4 years from a single center with a high MIC_90_ value for FLZ and AMB. Although our research was limited to examining past cases from one hospital and conducting tests in lab settings without linking the data to how patients were treated, it still offers valuable insights into *Candida* infections and their responses to different antifungal medications, which can aid in reducing illness and fatalities. The emergence of new and rare species further complicates therapeutic strategy due to limited data availability of ECVs and their breakpoints.

## ABBREVIATION

MALDI-TOF MS: Matrix-assisted laser desorption/ionization time-of-flight mass spectrometry
CLSI: Clinical and Laboratory Standards Institute
MIC: Minimum inhibitory concentration
NICU: Neonatal Intensive Care Unit
ECV: Epidemiological cutoff values

## AUTHOR CONTRIBUTIONS

Punit Tiwari investigated, performed formal analysis, methodology, data curation, and visualization, and drafted the manuscript. Munesh Kumar Gupta and Amit Kumar Rai performed an analysis of the data. All work was done under the supervision of Ragini Tilak and Pradip Paik, and all authors approved the manuscript.

## ACKNOWLEDGEMENTS

We would like to acknowledge for financial supports, Indian Council of Medical Research (ICMR), India (Ref: EMDR/SG/12/2023-4724), Anusandhan National Research Foundation (ANRF), India (Ref: CRG/2023/005576), CSIR-UGC New Delhi, India (JRF Grants no. 1055/(CSIR-UGC NET JUNE 2019) for Punit Tiwari, and Institute of Eminence (IoE), BHU, Varanasi, India, and Institute of Medical Sciences (IMS) BHU for supporting with patients samples, and experimental facilities.

## Ethical approval

This study received ethical clearance from the Institutional Review Board with Reference number ECR/526/Inst/UP/2019.

## DATA AVAILABILITY

All data generated during and/or analyzed during the current research project are available from the corresponding author on reasonable request.

## REFERENCES

1. Pfaller MA, Diekema DJ. 2007. Epidemiology of invasive candidiasis: A persistent public health problem. Clin Microbiol Rev 20:133–163. 10.1128/cmr.00029-06

2. Tak V, Mathur P, Varghese P, Gunjiyal J, Xess I, Misra MC. 2014. The Epidemiological Profile of Candidemia at an Indian Trauma Care Center. J Lab Physicians 6:096–101. 10.4103/0974-2727.141506

3. Giri S, Kindo AJ. 2012. A review of Candida species causing blood stream infection. Indian J Med Microbiol 30:270–278. 10.4103/0255-0857.99484

4. Bajwa S, Kulshrestha A. 2013. Fungal infections in intensive care unit: Challenges in diagnosis and management. Ann Med Health Sci Res 3:238. 10.4103/2141-9248.113669

5. Falagas ME, Roussos N, Vardakas KZ. 2010. Relative frequency of albicans and the various non-albicans Candida spp among candidemia isolates from inpatients in various parts of the world: A systematic review. Int J Infect Dis 14:e954–e966. 10.1016/j.ijid.2010.04.006

6. Falagas ME, Apostolou KE, Pappas VD. 2006. Attributable mortality of candidemia: A systematic review of matched cohort and case-control studies. Eur J Clin Microbiol Infect Dis 25:419–425. 10.1007/s10096-006-0159-2

7. Turner SA, Butler G. 2014. The Candida pathogenic species complex. Cold Spring Harb Perspect Med 4:1–17. 10.1101/cshperspect.a019778

8. Jiang C, Dong D, Yu B, Cai G, Wang X, Ji Y, Peng Y. 2013. Mechanisms of azole resistance in 52 clinical isolates of Candida tropicalis in China. J Antimicrob Chemother 68:778–785. 10.1093/jac/dks481

9. Yesudhason BL, Mohanram K. 2015. Candida tropicalis as a predominant isolate from clinical specimens and its antifungal susceptibility pattern in a tertiary care hospital in Southern India. J Clin Diagnostic Res 9:DC14–DC16. 10.7860/JCDR/2015/13460.6208

10. Dagi HT, Findik D, Senkeles C, Arslan U. 2016. Identification and antifungal susceptibility of Candida species isolated from bloodstream infections in Konya, Turkey. Ann Clin Microbiol Antimicrob 1–5. 10.1186/s12941-016-0153-1

11. Berkow EL, Lockhart SR, Ostrosky-zeichner L. 2020. Antifungal Susceptibility Testing : Current Approaches. ClinicalMicrobiologyReviews 33:1–30. 10.1128/cmr.00069-19

12. Arastehfar A, Daneshnia F, Kord M, Roudbary M. 2019. Comparison of 21-Plex PCR and API 20C AUX, MALDI-TOF MS, and rDNA Sequencing for a Wide Range of Clinically Isolated Yeast Species : Improved Identification by Combining 21-Plex PCR and API 20C AUX as an Alternative Strategy for Developing Countries. Front Cell Infect Microbiol 9:1–9. 10.3389/fcimb.2019.00021

13. Ghosh AK, Paul S, Sood P, Rudramurthy SM, Rajbanshi A, Jillwin TJ, Chakrabarti A. 2015. Matrix-assisted laser desorption ionization time-of-flight mass spectrometry for the rapid identification of yeasts causing bloodstream infections. Clin Microbiol Infect 21:372–378. 10.1016/j.cmi.2014.11.009

14. Posteraro B, Spanu T, Fiori B, De Maio F, De Carolis E, Giaquinto A, Prete V, De Angelis G, Torelli R, D’Inzeo T, Vella A, De Luca A, Tumbarello M, Ricciardi W, Sanguinetti M. 2015. Antifungal susceptibility profiles of bloodstream yeast isolates by sensititre yeastone over nine years at a large Italian teaching hospital. Antimicrob Agents Chemother 59:3944–3955. 10.1128/aac.00285-15

15. Fisher MC, Alastruey-Izquierdo A, Berman J, Bicanic T, Bignell EM, Bowyer P, Bromley M, Brüggemann R, Garber G, Cornely OA, Gurr SJ, Harrison TS, Kuijper E, Rhodes J, Sheppard DC, Warris A, White PL, Xu J, Zwaan B, Verweij PE. 2022. Tackling the emerging threat of antifungal resistance to human health. Nat Rev Microbiol 20:557–571. 10.1038/s41579-022-00720-1

16. Alexander BD. 2012. Document M27-A4. Ref Method Broth Dilution Antifung Suscetibility Test Yeasts.

17. CLSI. 2026. Performance standards for antifungal susceptibility testing of yeasts. In CLSI supplement M27M44S, 4th ed. Clinical and Laboratory Standards Institute.

18. Mckinney W. 2010. Data Structures for Statistical Computing in Python. scipy 1:56–61.

19. Pedregosa F, Weiss R, Brucher M. 2011. Scikit-learn : Machine Learning in Python. J Mach Learn Res 12:2825–2830.

20. Zeng G. 2023. On the analytical properties of category encodings in logistic regression. Commun Stat - Theory Methods 0:1–18. 10.1080/03610926.2021.1939382

21. Breiman L. 2001. Random forests. Mach Learn 45:5–32.

22. Stone M. 1974. Cross-Validatory Choice and Assessment of Statistical Predictions. J R Stat Soc Ser B 36:111–147. 10.1111/j.2517-6161.1974.tb00994.x

23. Sokolova M, Lapalme G. 2009. A systematic analysis of performance measures for classification tasks. Inf Process Manag 45:427–437. 10.1016/j.ipm.2009.03.002

24. Puig-Asensio M, Ruiz-Camps I, Fernández-Ruiz M, Aguado JM, Muñoz P, Valerio M, Delgado-Iribarren A, Merino P, Bereciartua E, Fortún J, Cuenca-Estrella M. 2015. Epidemiology and outcome of candidaemia in patients with oncological and haematological malignancies: Results from a population-based surveillance in Spain. Clin Microbiol Infect 21:491.e1–491.e10. 10.1016/j.cmi.2014.12.027

25. Pandey N, Gupta MK, Paul P, Tilak R. 2020. Necessity to identify candida species accurately with minimum inhibitory concentration determination in each case of bloodstream infections. J Infect Public Health 13:753–758. 10.1016/j.jiph.2019.12.002

26. Lockhart SR, Iqbal N, Cleveland AA, Farley MM, Harrison LH, Bolden CB, Baughman W, Stein B, Hollick R, Park BJ, Chiller T. 2012. Species identification and antifungal susceptibility testing of Candida bloodstream isolates from population-based surveillance studies in two U.S. cities from 2008 to 2011. J Clin Microbiol 50:3435–3442. 10.1128/jcm.01283-12

27. Otto C, Babady NE. 2023. Epidemiology and Outcomes of Non-albicans Candida Bloodstream Infections in Transplant Recipients and Cancer Patients. Mycopathologia 188:863–871. 10.1007/s11046-023-00765-7

28. Marins, et al. 2018. Evaluation of Candida bloodstream infection and antifungal utilization in a tertiary care hospital. BMC Infect Dis 18:1–13. 10.1186/s12879-018-3094-9

29. Lukic-Grlić A, Mlinarić-Missoni E, Škaric I, Važic-Babić V, Svetec IK. 2011. Candida utilis candidaemia in neonatal patients. J Med Microbiol 60:838–841. 10.1099/jmm.0.023408-0

30. Shivadasan J, Raksha K, Urs PS. 2016. Candida utilis causing neonatal Candidemia – A case report and literature review. Apollo Med 13:55–58. 10.1016/j.apme.2016.01.001

31. Rai AK, Tilak R, Tiwari P, Meena P, Kumar A, Tiwari AK, Gupta MK. 2024. Original article Emergence of rare and uncommon yeast-like pathogens causing neonatal sepsis at a tertiary care center, North India. Germs 14:20–27. 10.18683/germs.2024.1414

32. Neves RP, Parahym AMR de C, Silva CM da, Macêdo DPC, Leal AFG, Neves HJ, Lima-Neto RG. 2018. Fungal Infections in Neonatal Intensive Care. Sel Top Neonatal Care 10.5772/intechopen.70302.

33. Nami S, Aghebati-Maleki A, Morovati H, Aghebati-Maleki L. 2019. Current antifungal drugs and immunotherapeutic approaches as promising strategies to treatment of fungal diseases. Biomed Pharmacother 110:857–868. 10.1016/j.biopha.2018.12.009

34. Fothergill AW, Sutton DA, McCarthy DI, Wiederhold NP. 2014. Impact of new antifungal breakpoints on antifungal resistance in Candida species. J Clin Microbiol 52:994–997. 10.1128/jcm.03044-13

35. Mokaddas EM, Al-Sweih NA, Khan ZU. 2007. Species distribution and antifungal susceptibility of Candida bloodstream isolates in Kuwait: A 10-year study. J Med Microbiol 56:255–259. 10.1099/jmm.0.46817-0

36. Jain K, Shukla N, Kaur A, Kalita B, Srivastava S, Yadav S, Rani P, Kagra P, Singh A, Chaudhary K, Kaur H, Patel J, Chauhan M, Tarai B, Narayan J. 2026. Genomic insights into a clade-specific Candida tropicalis lineage with resistance to azoles and immune evasion traits. MBio 17:1–23. 10.1128/mbio.00235-26

37. Misas E, Witt LS, Farley MM, Thomas S, Jenkins EN, Gade L, Peterson JG, Restrepo AM, Fridkin S, Lockhart SR, Chow NA, Lyman M. 2024. Molecular and Epidemiological Investigation of Fluconazole-resistant Candida parapsilosis - Georgia, United States, 2021. Open Forum Infect Dis 11:1–9. 10.1093/ofid/ofae264

38. Pfaller MA, Diekema DJ, Gibbs DL, Newell VA, Ellis D, Tullio V, Rodloff A, Fu W, Ling TA. 2010. Results from the artemis disk global antifungal surveillance study, 1997 to 2007: A 10.5-year analysis of susceptibilities of candida species to fluconazole and voriconazole as determined by CLSI standardized disk diffusion. J Clin Microbiol 48:1366–1377. 10.1128/jcm.02117-09

39. Wisplinghoff H, Ebbers J, Geurtz L, Stefanik D, Major Y, Edmond MB, Wenzel RP, Seifert H. 2014. Nosocomial bloodstream infections due to Candida spp. in the USA: Species distribution, clinical features and antifungal susceptibilities. Int J Antimicrob Agents 43:78–81. 10.1016/j.ijantimicag.2013.09.005

40. Fothergill AW, Sutton DA, Mccarthy DI, Wiederhold NP. 2014. Impact of New Antifungal Breakpoints on Antifungal Resistance in Candida Species. J Clin Microbiol 52:994–997. 10.1128/jcm.03044-13

41. Bilal H, Shafiq M, Hou B, Islam R, Nadeem M, Ullah R. 2022. Distribution and antifungal susceptibility pattern of Candida species from mainland China : A systematic analysis. Virulence 13:1573–1589. 10.1080/21505594.2022.2123325

42. Kumar S, Kumar A, Roudbary M, Mohammadi R, Černáková L, Rodrigues CF. 2022. Overview on the Infections Related to Rare Candida Species. Pathogens 11:1–45. 10.3390/pathogens11090963

43. Xie Y, Peng Q, Ji Y, Xie A, Yang L, Mu S, Li Z, He T, Xiao Y, Zhao J, Zhang Q. 2021. Isolation and Identification of Antibacterial Bioactive Compounds From Bacillus megaterium L2. Front Microbiol 12:1–10. 10.3389/fmicb.2021.645484

44. Sreelekshmi TS, Ninan MM, Premanand A, Chacko A, Sahni RD, Michael JS. 2021. Candida utilis : a rare cause of septicemia in children 3–6. 10.1099/acmi.0.000281

45. Pfaller MA, Moet GJ, Messer SA, Jones RN, Castanheira M. 2011. Candida bloodstream infections: Comparison of species distributions and antifungal resistance patterns in community-onset and nosocomial isolates in the SENTRY Antimicrobial Surveillance Program, 2008-2009. Antimicrob Agents Chemother 55:561–566. 10.1128/aac.01079-10

46. Salama EA, Elgammal Y, Utturkar SM, Lanman NA, Hazbun TR, Seleem MN. 2024. Overcoming amphotericin B resistance in Candida auris using the antiemetic drug rolapitant. Antimicrob Agents Chemother 68. 10.1128/aac.00556-24

47. Cavassin FB, Baú-Carneiro JL, Vilas-Boas RR, Queiroz-Telles F. 2021. Sixty years of Amphotericin B: An Overview of the Main Antifungal Agent Used to Treat Invasive Fungal Infections. Infect Dis Ther 10:115–147. 10.1007/s40121-020-00382-7

48. Mesa-Arango AC, Scorzoni L, Zaragoza O. 2012. It only takes one to do many jobs: Amphotericin B as antifungal and immunomodulatory drug. Front Microbiol 3:1–10. 10.3389/fmicb.2012.00286

49. Pfaller MA, Diekema DJ, Gibbs DL, Newell VA, Nagy E, Dobiasova S, Rinaldi M, Barton R, Veselov A, Surveillance A. 2008. Candida krusei, a Multidrug-Resistant Opportunistic Fungal Pathogen : Geographic and Temporal Trends from the ARTEMIS DISK Antifungal Surveillance Program, 2001 to 2005. J Clin Microbiol 46:515–521. 10.1128/jcm.01915-07

50. Kathuria S, Singh PK, Sharma C, Prakash A, Masih A, Kumar A, Meis JF, Chowdhary A. 2015. Multidrug-resistant Candida auris misidentified as Candida haemulonii: Characterization by matrix-assisted laser desorption ionization-time of flight mass spectrometry and DNA sequencing and its antifungal susceptibility profile variability by vitek 2, CLSI Broth Microdilution, and Etest Method. J Clin Microbiol 53:1823–1830. 10.1128/jcm.00367-15

51. Monteiro AÉJ, Sidrim JOSÉJC, Rocha MFG. 2013. Minimum inhibitory concentrations of amphotericin B, azoles and caspofungin against Candida species are reduced by farnesol. Med Mycol 53–59. 10.3109/13693786.2012.692489

52. Solomon MP, Mahler B, Popescu O, Ditu L, Gheorghe-barbu I, Chivu LI, Sabina L, Manolescu C. 2026. Shifting Epidemiology and Antifungal Susceptibility Patterns of Clinical Fungal Isolates in an Intensive Care Unit (ICU) from Bucharest, Romania : A Retrospective Observational Study 1–21. 10.3390/antibiotics15050440

53. Vahedi-shahandashti R, Lass-flörl C. 2025. In vitro activity of SF001: a next-generation polyene versus amphotericin B. Antimicrob Agents Chemother 1–5. 10.1128/aac.00322-25

54. Pfaller MA, Diekema DJ, Rex JH, Alexander BD, Andes D, Brown SD, Chaturvedi V, Ghannoum MA, Knapp CC, Sheehan DJ, Walsh TJ. 2008. Correlation of MIC with Outcome for Candida Species Tested against Caspofungin, Anidulafungin, and Micafungin : Analysis and Proposal for Interpretive MIC Breakpoints. J Clin Microbiol 46:2620–2629. 10.1128/jcm.00566-08

55. Pfaller MA, Diekema DJ, Andes D, Arendrup MC, Brown SD, Lockhart SR, Motyl M, Perlin DS, Testing A. 2011. Clinical breakpoints for the echinocandins and Candida revisited : Integration of molecular, clinical, and microbiological data to arrive at species-specific interpretive criteria. Drug Resist Updat 14:164–176. 10.1128/cmr.00029-06

